# Genome-wide patterns of local adaptation in *Drosophila melanogaster*: adding intra European variability to the map

**DOI:** 10.1101/269332

**Authors:** Lidia Mateo, Gabriel E. Rech, Josefa González

## Abstract

Signatures of spatially varying selection have been investigated both at the genomic and transcriptomic level in several organisms. In *Drosophila melanogaster*, the majority of these studies have analyzed North American and Australian populations, leading to the identification of several loci and traits under selection. However, populations in these two continents showed evidence of admixture that likely contributed to the observed population differentiation patterns. Thus, disentangling demography from selection is challenging when analyzing these populations. European populations could be a suitable system to identify loci under spatially varying selection provided that no recent admixture from African populations would have occurred. In this work, we individually sequence the genome of 42 European strains collected in populations from contrasting environments: Stockholm (Sweden), and Castellana Grotte, (Southern Italy). We found low levels of population structure and no evidence of recent African admixture in these two populations. We thus look for patterns of spatially varying selection affecting individual genes and gene sets. Besides single nucleotide polymorphisms, we also investigate the role of transposable elements in local adaptation. We concluded that European populations are a good dataset to identify loci under spatially varying selection. The analysis of the two populations sequenced in this work in the context of all the available *D. melanogaster* data allowed us to pinpoint genes and biological processes relevant for local adaptation. Identifying and analyzing populations with low levels of population structure and admixture should help to disentangle selective from non-selective forces underlying patterns of population differentiation in other species as well.

## INTRODUCTION

Adaptation is a central concept in evolutionary biology. Adaptation underlies such important processes as the ability of species to survive in changing environments. However, understanding how organisms adapt to their environment, specifically which genes and which biological processes are more relevant for adaptation, are still open questions in evolutionary biology. In the last decades, adaptation to new environments associated with colonization processes, and adaptation to changing environmental conditions, both natural and human-driven, has been extensively studied (reviewed in Adrion *et al.* (2015); Fan *et al.* (2016) and Flood & Hancock (2017)). Analyses of several populations collected along environmental gradients, and of pairs of populations from contrasting environments are often used to identify genetic variants under spatially varying selection. Several approaches have been applied to the detection of adaptive evolution, from investigating the genetic basis of a known adaptive trait, to the identification of candidate adaptive variants without *a priori* knowledge of the relevant adaptive phenotypes (Pardo-Diaz *et al.* 2015; Hoban *et al.* 2016). All these approaches have been boosted by the availability and reduced costs of sequencing technologies that are being applied at the genomic and transcriptomic level (Pardo-Diaz *et al.* 2015; Villanueva-Cañas *et al.* 2017). Insights on the genomic targets of spatially varying selection, the molecular mechanisms underpinning local adaptation, and the extent of parallel adaptation, in model and non-model species, is starting to accumulate (Adrion *et al.* 2015; Fan *et al.* 2016 and Flood & Hancock 2017). However, several challenges such as the role of demography in the observed population differentiation patterns, and thus the identification of the true targets of spatially varying selection (Flatt 2016) still remain. In this work, we tested whether European *Drosophila melanogaster* natural populations could be a good model system to disentangle the effects of demography and selection, and thus to identify the genetic basis of local adaptation.

*D. melanogaster* has several characteristics that makes it an exceptional organism to pinpoint adaptive variants and thus to advance our knowledge on the genes and biological processes that are more relevant for adaptation. This species has recently (10.000-16.000 years ago) colonized worldwide environments from its ancestral range in subtropical Africa (Li & Stephan 2006). This colonization required adaptation to both biotic and abiotic factors present in the new environments. Thus a wide-range of adaptations should be common in this species, and they should still be detectable as partial or complete sweeps at the DNA level (Przeworski 2002). Moreover, in *D. melanogaster* the identified genetic variants can be mapped to a well-annotated genome, and the wealth of genetic tools available allows to experimentally test the molecular mechanism and fitness consequences of the identified genetic variants (Mohr *et al.* 2014). Not surprisingly, *D. melanogaster* has been extensively used as a model organism to identify the genetic targets of spatially varying selection (Hoffmann & Weeks 2007; Adrion *et al.* 2015). These studies have identified genetic variants and traits involved in geographical adaptation although only rarely genetic variants have been connected to fitness-related traits (Paaby & Schmidt 2008; Schmidt *et al.* 2008; Lee *et al.* 2011; Magwire *et al.* 2012; Lee *et al.* 2013; Guio *et al.* 2014; Mateo *et al.* 2014; Paaby *et al.* 2014; Ullastres *et al.* 2015; Merenciano *et al.* 2016).

Most of our knowledge on the genetic basis of spatially varying selection so far comes from the analyses of North American and Australian *D. melanogaster* populations (reviewed in Adrion *et al.* 2015). However, the identification of genes underlying local adaptation in populations from these two continents is confounded by the admixture between European and African populations (Caracristi & Schlotterer 2003; Yukilevich & True 2008b, a; Yukilevich *et al.* 2010; Duchen *et al.* 2013; Fabian *et al.* 2015; Kao *et al.* 2015; Bergland *et al.* 2016). In both continents, the secondary contact between *D. melanogaster* populations from the derived and ancestral ranges of the species creates population differentiation patterns similar to those created by spatially varying selection. Thus, identification of clinally variant loci cannot be taken as evidence of spatially varying selection without considering also the potential role of admixture in the patterns observed (Ullastres *et al.* 2015; Bergland *et al.* 2016). Looking for overlapping patterns of population differentiation in closely related species has recently been used to overcome this limitation (Zhao *et al.* 2015; Machado *et al.* 2016). Comparing the stability of clines over long time periods should also help distinguish between polymorphisms maintained by selection and polymorphisms resulting from recent admixture (Anderson *et al.* 2005; Umina *et al.* 2005; Weeks *et al.* 2006; Cogni *et al.* 2014; Kapun *et al.* 2016; Cogni *et al.* 2017). Alternatively, comparing populations from other continents in which recent admixture has not occurred could also help in the identification of loci under spatially varying selection. There are a few studies in which European populations have been analyzed (Hutter *et al.* 2008; Muller *et al.* 2011; Catalan *et al.* 2012; Bozicevic *et al.* 2016). However, they were all based on the comparison between European and African strains, and to our knowledge, there is no study in which patterns of population differentiation within the European continent have been analyzed. Thus, whether European populations do not show recent admixture and as such are a good system to identify loci under spatially varying selection remains to be tested. Another characteristic of the studies performed to date is that most of them are based on the analysis of a single type of genetic variant, which most often is Single Nucleotide Polymorphisms (SNPs). Other type of variants that are also likely to play a role in adaptation, such as transposable element (TE) insertions, other copy number variants, and inversions, remain understudied (Gonzalez *et al.* 2008, 2010, 2015; Kapun *et al.* 2016; Schrider *et al.* 2016). TEs are particularly likely to be involved in adaptation because they generate mutations that often have complex phenotypic effects (Chuong *et al.* 2017; Horvath *et al.* 2017). In a series of analyses, Gonzalez *et al.* (2008, 2010, 2015), investigated a subset of TE insertions in two North American, two Australian, and two populations collected in opposite slopes of the Evolution Canyon, in Israel, that although being geographically close have temperate- and tropical-like climates. While some insertions showed parallel patterns in North America and Australia, none of the investigated TEs differed in the Evolution Canyon populations suggesting that they might not be targets of spatially varying selection (González *et al.* 2015). However, the dataset of TEs analyzed in these studies was incomplete and biased. Thus, further analyses are needed to get a more complete picture of the role of TEs in local adaptation (González *et al.* 2015).

In this work, we aimed at testing whether European *D. melanogaster* populations could be a good model system to disentangle demography from selection, and thus to identify genomic targets of spatially varying selection. To accomplish this, we first tested whether two European populations collected in localities with contrasting environments: Stockholm in Sweden, which has a humid continental climate, and Castellana Grotte in Bari, Southern Italy, which has a hot summer Mediterranean climate, showed evidence of population structure or admixture patterns (Figure 1). These analyses were performed in the context of a more global dataset of natural populations including one African population from the ancestral range of the species (Gikongoro, Rwanda, Pool *et al.* (2012)) and one North American population (Raleigh, North Carolina, Huang *et al.* (2014)) (Figure 1, Table S1, Supporting information). After excluding a major role of population structure and admixture in the genomic differentiation patterns of these two European populations, we used F_ST_ to identify both SNPs and TE insertions under selection. In addition to identifying individual candidate genes, we also performed gene set enrichment analyses, which allowed us to identify more subtle trends in allele frequency changes affecting several genes that are indicative of polygenic adaptation (Daub *et al.* 2013). Finally, we draw on several independent sources of evidence to identify a set of genomic targets of spatially varying selection in *D. melanogaster* natural populations.

**Figure 1.**
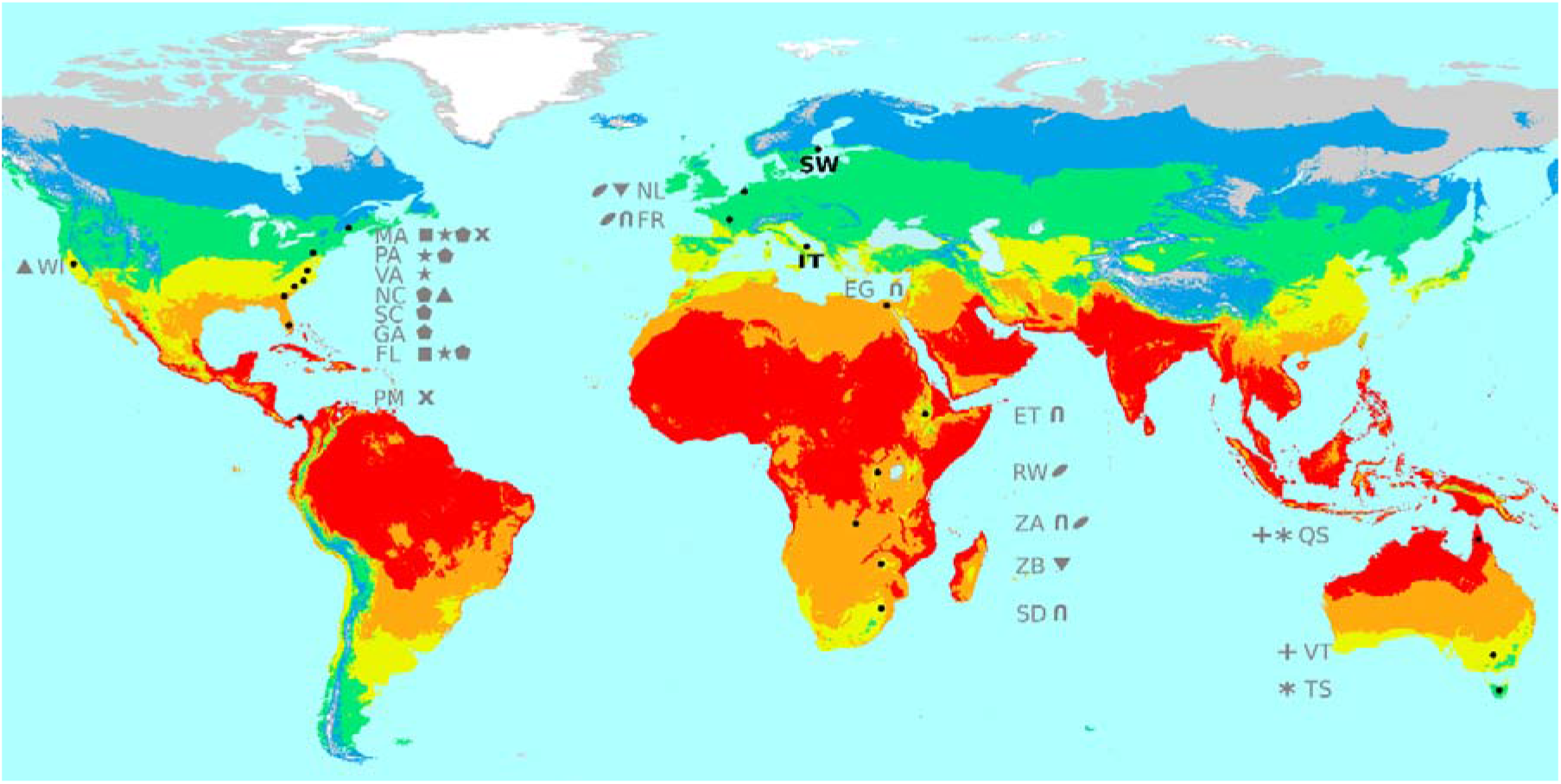
Geographical location of the two European populations analyzed in this work. SW (Sweden) and IT (Italy) populations are showed in bold. In addition, locations from where natural populations were analyzed looking for differentiated genes are showed in grey: FL: Florida, MA: Maine, TS: Tasmania, QS: Queensland, GA: Georgia, NC: North Carolina, SC: South Carolina, PA: Pennsylvania, WI: Winters, VA: Virginia, FR: France, EG: Egypt, ET: Ethiopia, SD: South Africa, ZA: Zambia, VT: Victoria, PM: Panama, ZB: Zimbabwe, NL: Netherlands, RW: Rwanda. Symbols indicate populations used in the same study (See Table S1, Supporting information).

## MATERIAL AND METHODS

### Sample collection and newly sequenced genomes

Genomes reported here are derived from two natural population samples collected in Stockholm (59.34°, 17.94°, Sweden) and in Castellana Grotte (40.90°, 17.16°, Bari, Southern Italy) in September and October 2011, respectively (Figure 1 and Table S2, Supporting information). Isofemale lines were established from a single wild-caught female and maintained in the laboratory for at least twenty generations at room temperature before sequencing. A total of 26 Swedish and 16 Italian strains were individually sequenced. For each strain, DNA was extracted from 30 adult females using Puregene Core Kit A (Qiagen, Germany) according to the manufacturer’s instructions. Short-sequence libraries (∼500bp) were constructed and paired-end sequencing using Illumina High-Seq2000 platform was performed (100bp reads). Quality control was performed using FastQC (Andrews 2010). Raw fastq files were deposited in the Sequence Read Archive (SRA) from the National Center for Biotechnology Information (NCBI) under the BioProject accession: PRJNA390275.

### Previously sequenced genomes

A total of 24 inbred strains collected in Raleigh (North Carolina, USA) from the DGRP panel (Mackay *et al.* 2012; Huang *et al.* 2014), and haploid embryos derived from 17 inbred or isofemale strains collected in Gikongoro (Rwanda, Africa) from the DPGP2 panel (Pool *et al.* 2012) were also analyzed (Table S2, Supporting information). All these strains were sequenced to a similar depth and quality as the European strains reported in this work (Table S2, Supporting information). The 17 Rwandan strains are a subset of the 22 strains available for this population with no inferred cosmopolitan admixture (nine strains) or < 3% admixture (eight strains). For all these strains, raw reads in *fastq* format were downloaded from the NCBI-SRA. The global cohort consisted of a total of 83 genomes (26 Swedish, 16 Italian, 24 North American, and 17 African) that were analyzed using the same pipeline (Table S2, Supporting information).

### Read mapping and coverage analysis

Reads from the four populations analyzed were quality trimmed using *sickle v.1.2* when mean quality dropped below 20 (Joshi & Fass 2011). Reads shorter than 50bp were discarded. Read mapping to the *D. melanogaster* reference genome (Flybase release 5) was performed using *bwa-Backtrack v. 0.78* algorithm using strict default parameters (Li & Durbin 2010). Optical duplicates were marked with *Picard v.1.115.* The alignment around indels was refined using *GATK v.3.2 IndelRealigner* (McKenna *et al.* 2010). Reads with a BWA mapping quality score less than 20 and unmapped reads were realigned using *Stampy v.1.0.20* for a more accurate re-mapping (Lunter & Goodson 2011). Reads with a *Stampy* mapping score higher than 20 were merged with the previous alignment. The distribution of coverage per base pair was calculated for each strain using *GATK BaseCoverageDistribution.* The coverage values corresponding to the 0.975 percentile were calculated for each strain using a custom R script (Table S2, Supporting information).

### Variant Calling and Joint Genotyping

We followed the two-step GATK variant calling pipeline as follows. First, haplotype calling was performed independently for each strain using *GATK HaplotypeCaller* with a minimum base quality of 31 (Mackay *et al.* 2012). We checked the distribution of heterozygous SNPs and they were distributed along the genome without any apparent region being enriched for heterozygous SNPs. Second, joint genotyping of the cohort of 83 genomes was performed using *GATK GenotypeGVCF,* which aggregates the gVCF files obtained from *GATK HaplotypeCaller* into a multi-sample VCF file. A total of 6,820,800 variants were called, including indels. Out of 6,820,800 variants, 5,479,546 were annotated as SNPs. SNPs falling in low-recombination regions close to centromeres and telomeres were excluded. The coordinates corresponding to euchromatic regions used in our analyses were defined by the recombination rate calculator (RRC) estimates (Fiston-Lavier *et al.* 2010) and included 2L:530,000–18,870,000; 2R:1,870,000–20,860,000; 3L:750,000–19,020,000; 3R:2,580,000–27,440,000; and X:1,220,000–21,210,000. These regions contained a total of 4,804,172 SNPs. SNPs within 5 bp of indels or other types of complex mutations and SNPs falling in repetitive or low complexity regions according to RepeatMasker were excluded to obtain a set of 4,106,958 SNPs. Different further filters were specifically applied in each analysis and are detailed in the corresponding sections (see below).

### Genetic diversity and population structure analysis in the four worldwide populations

Regions of known cosmopolitan admixture in the African strains were excluded from the initial dataset of 4,106,958 SNPs, yielding a total of 3,749,072 SNPs (Pool *et al.* 2012). Genome-wide diversity levels (π) were estimated for non-overlapping windows of 10,000 SNPs using *vcftools v.0.1.12*. The average nucleotide diversity was calculated for the whole genome and separately for the autosomes and the X chromosome using an *ad hoc* R script.

We calculated nucleotide diversity considering that we were sampling two alleles. Because it is possible that in each isofemale strain genome we are sampling greater than or less than two alleles per locus, we also estimated nucleotide diversity after sampling one allele per genome with a weighted probability based on allelic read counts using an *ad hoc* python script.

Global weighted *Weir and Cockerham* unbiased estimator of the fixation index (F_ST_) was estimated between each pair of populations using *vcftools* (Weir & Cockerham 1984; Danecek *et al.* 2011). Population structure was also assessed by principal component analysis using *Eigensoft* and model-based global ancestry estimation using *Admixture*. SNPs with a minor allele frequency <15% or a genotyping rate lower than 70% were excluded. Two African strains with a variant calling lower than 70% (RG4, RG18) were also excluded. The final set consisted on 68,390 high confidence SNPs. Model-based global ancestry was estimated using *Admixture* software (Alexander *et al.* 2009). The number of ancestral populations analyzed (*k*) ranged from 1 to 7. Cross validation error for each value of *k* was used to determine the most likely hypothesis. The ancestry coefficients for each strain were plotted using the *ggplot2 v1.0.0* library loaded in *R v-3.1.0*.

### Testing for population differentiation in Europe

The following additional filters were applied to the initial set of 4,106,958 SNPs: (i) only non-singleton biallelic SNPs were considered; (ii) A custom perl script was used to filter to *N* individual calls with coverage lower than 10x or higher than the 0.975 percentile of the coverage per base pair distribution for the strain considered (see Table S2, Supporting information); (iii) SNPs with a genotyping rate lower than 70% were excluded. In our dataset, this threshold maximizes the number of positions that pass the filtering while excluding those positions with an excess of missing genotypes. (iv) SNPs segregating at a frequency lower than 5% in the European population were excluded.

A total of 1,123,227 SNPs passed our conservative filtering pipeline in the European populations and were used to calculate the pairwise *Weir and Cockerham* unbiased estimator of the fixation index (F_ST_) using *vcftools* (Weir & Cockerham 1984; Danecek *et al.* 2011). Significance of the heterogeneity in population differentiation levels across chromosomal arms was determined by performing one-way ANOVA. In subsequent analyses, the empirical distribution of F_ST_ in each chromosomal arm was used to identify outliers. Genomic differentiation within the In(2L)t common cosmopolitan inversion (2L:2225744..13154180, (Pool *et al.* 2012)) was compared to the levels observed in the rest of the chromosomal arm by performing a Mann-Whitney U test. We also estimated the population frequency of common and rare cosmopolitan inversions in the 42 European strains analysed in this work using the panel of SNPs known to be associated with these inversions (Kapun *et al.* 2016).

### Over/Under representation of candidate differentiated loci per functional category

Functional annotation of SNPs was performed using *VariantAnnotation* R/Bioconductor package (Obenchain *et al.* 2014). Out of the initial 1,123,227 SNPs, 1,121,948 could be annotated into the following functional categories: intergenic, promoter (1,000 bp upstream of the transcription start site), core promoter (316 bp upstream of the transcription start site; (Hoskins *et al.* 2011), 5’ UTR, coding, splice site, intron, and 3’ UTR using *TxDb.Dmelanogaster.UCSC.dm3.ensGene* R/Bioconductor package *v.2.14.0*. Coding variants were further classified into synonymous and non-synonymous. In order to obtain a set of neutrally evolving polymorphisms, we extracted those SNPs falling within the first 8-30 base pairs of small introns (<=65 bp) (Parsch *et al.* 2010). The rest of intronic SNPs, which mostly belong to large introns, were classified as non-small intronic SNPs.

Counts of SNPs falling in the top 5%, 1% and 0.5% tails of the chromosomal empirical F_ST_ distribution were determined for each functional category. Two sided Fisher’s exact test was used to determine if there were non-random associations between functional category and belonging to the 95% of the chromosomal arm F_ST_ distribution, or to the top 5%, 1% or 0.5% tails, while controlling for allele frequency, chromosomal arm, and considering SNPs in small introns as background of our analyses. The same procedure was used to determine if there was an over- or under-representation of candidate differentiated loci specifically located inside In(2L)t inversion with respect to the rest of each chromosomal arm.

### Assignment of Z_ST_ scores to genes

A population differentiation score per gene was obtained while controlling for gene length bias. SNP-wise F_ST_ values were converted to gene-wise F_ST_ values by assigning to each gene the maximum F_ST_ value among those SNPs found along its transcribed region and the 1kb region upstream of the transcription start site. This has been shown to be a good representation of the F_ST_ per gene (Daub *et al.* 2013). Maximum F_ST_ values per gene were normalized to a Z score using chromosome specific empirical F_ST_ distributions. There was a positive correlation between gene length (and number of SNPs) and Z score (r = 0.37132; p-value = 2.2e-16) (see Figure S1A, Supporting information). In order to correct for this potential bias, genes were assigned to 16 bins containing all genes with a similar number of SNPs and showing non-significant correlation between number of SNPs and Z score within each bin. We followed the approach detailed in (Daub *et al.* 2013) to obtain a median based corrected Z score that will be referred to as Z_ST_ score. As a result, the correlation between the number of SNPs per gene and Z_ST_ was eliminated (r = 0.01121; p-value = 0.1981) (Figure S1B, Supporting information). A total of 657 genes in the top 5% tail of the empirical Z_ST_ distribution were considered as candidate differentiated genes.

### Functional enrichment analysis

Gene Ontology (GO) term enrichment analysis in the list of 657 candidate differentiated genes was performed using *topGO v.2.16.0* (Alexa *et al.* 2006). We also performed gene set enrichment analysis by applying the Kolmogorov-Smirnov like test, which computes enrichment based on gene Z_ST_ scores using GO’s biological process annotation as gene set definition. In addition to the conventional Fisher’s exact test or the Kolmogorov-Smirnov test (*classic)*, we used three other algorithms: *elim, weight* and *weight01* to identify significantly enriched GO biological processes. While in the *classic* enrichment test each GO term is tested independently, in the other three algorithms the hierarchical structure of the GO is taken into account so that the enrichment score incorporates information about the whole GO topology. This eliminates local similarities and dependencies between GO terms and reduces the high false-positive rate that is usually attained with classic GO enrichment tests (Alexa *et al.* 2006). The *elim* method iteratively removes genes annotated to significant GO terms from their ancestors, following a bottom-up approach. This allows for the identification of more specific nodes in the GO graph. The *Weight* algorithm identifies the locally most significant terms in the GO graph (and not necessarily the most specific terms). Weight01 is a combination of *weight* and *elim* algorithms. We decided to look at the results yielded by *elim, weight* and *weight01* in order to explore which areas in the GO graph contain enriched GO terms instead of finding the precise GO terms that are enriched. Only GO terms with 5 or more differentiated genes were considered. We considered that a GO term was significantly enriched when the p-value computed by the *elim, weight* or *weight01* algorithms was smaller than 0.05. These methods are internally reducing the significance of most GO terms and practically disregarding many of them, reducing the probability of having false positives due to the multiple-testing problem. As such, the raw p-values obtained with this methods can be considered as corrected p-values (Alexa *et al.* 2006).

### TE frequency estimations

We used T-lex version 2 (*Tlex2*) to estimate presence/absence of TEs in each individual strain from the Italian and Sweden population (Fiston-Lavier *et al.* 2015). We used release 5 of the *Drosophila melanogaster* genome as reference along with TE coordinates downloaded from FlyBase (Hoskins *et al.* 2007). From the original 5,416 TE dataset, we excluded TEs overlapping other TEs, TEs that are part of segmental duplications, and TEs flanked by other repetitive regions (Fiston-Lavier *et al.* 2015). In addition, we excluded TEs belonging to the INE family (2,235 TEs) because we expect them to be fixed (Kapitonov & Jurka 2003; Singh & Petrov 2004; Yang & Barbash 2008). Overall, *Tlex2* was run for 1,632 TEs.

### F_ST_ calculation for TEs

Using the presence/absence information provided by *Tlex2* we created a multi-sample VCF file encoding the genotype of the 1,632 TEs in each individual strain. Then, we used *vcftools* (Danecek *et al.* 2011) for calculating the Weir & Cockerham (1984) estimator of the fixation index. A total of 556 TEs were at a frequency lower than 5% in both populations, 893 TEs were fixed in both populations, and for 22 TEs *Tlex2* was not able to determine the population frequencies. We ended up with F_ST_ values for 161 TEs. We compared the FST values obtained for the TEs to the empirical distribution of FST values obtained for SNPs. Those TEs falling in the top 5%, 2.5% or 0.5% tails of the distribution were considered as candidate differentiated TEs.

## RESULTS

### Genetic diversity, population structure, and admixture patterns in Swedish and Italian populations

We individually sequenced 26 strains collected in Stockholm, Sweden, and 16 strains collected in Castellana Grotte (Bari), Southern Italy, to an average coverage of 28.6x using Illumina technology (Figure 1 and Table S2, Supporting information).

To investigate the genetic diversity, population structure, and admixture patterns in European populations in the context of a more global dataset of natural populations, we used a joint genotyping strategy to call SNPs in strains from these two European populations, one North American population (24 strains collected in Raleigh, North Carolina, USA, Huang *et al.* (2014)), and one African population collected in the ancestral range of the species (17 strains sequenced from haploid embryos, Gikongoro, Rwanda, Africa, Pool *et al.* (2012)) (Table S2, Supporting information).

We estimated genome-wide diversity levels (π) for the two European populations sequenced in this study, and for the North American and African strains (Pool *et al.* 2012; Huang *et al.* 2014) (Figure 2 and Table S3a, Supporting information). The average level of nucleotide diversity is similar in the two European populations analyzed: 0.0047 and 0.0044 in Sweden and Italy, respectively. These values are similar to previous available estimates for two other European populations: Lyon (France) 0.0047 and Houten (Netherlands) 0.0046 (Lack *et al.* 2016). We also estimated genome-wide diversity levels sampling one allele per genome (see Material and Methods). In this case, estimates of genome-wide diversity were smaller (Table S3b, Figure S2, Supporting information).

**Figure 2.**
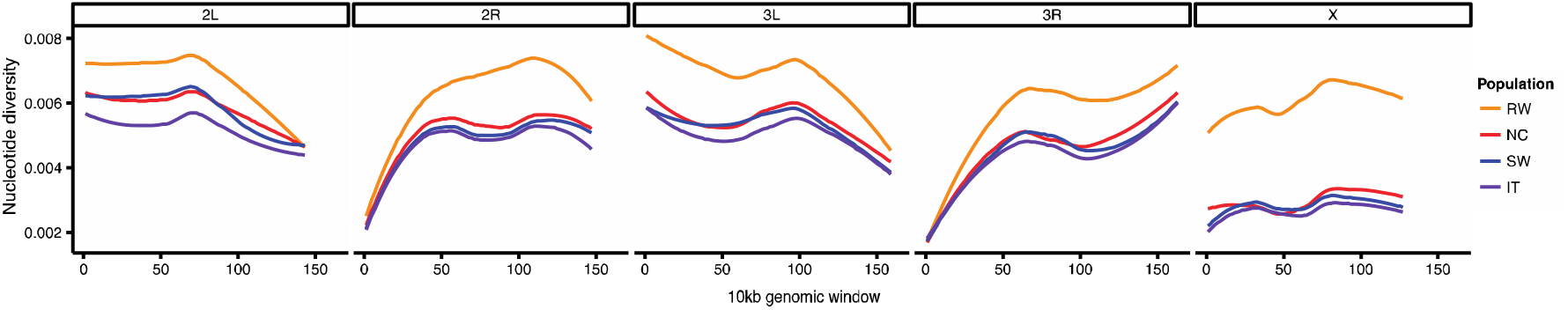
Genome-wide polymorphism (π) for each chromosomal arm. π estimates in 10.000 SNPs windows are plotted for each chromosome arm for each one of the four populations analyzed in this study.

Because the effective population size of the X chromosome is ¾ of that of the autosomes, we expect reduced levels of polymorphisms on the X chromosome compared to the autosomes (Begun and Whitley 1997). Indeed, we found that the average polymorphism on the X chromosome is reduced relative to autosomes both in the Swedish and Italian populations (Figure 2 and Table S3, Supporting information). On the other hand, we did not find reduced polymorphism in the X vs the autosomes in Rwanda (Figure 2 and Table S3, Supporting information). This has previously been reported, and it is likely due to an unequal sex ratio in African populations, where female population size has been estimated to be 1.8 times larger than male population size (Hutter *et al.* 2007). According to the African origin of *D. melanogaster,* diversity in non-African populations is expected to be a subset of that within Africa (David and Capy 1988). Consistently, we found that the Rwanda population showed the highest genomic diversity of the four populations analyzed (Figure 2).

Global F_ST_ values showed that the European populations are the less differentiated (Figure 3). F_ST_ between African and European populations is higher than between African and the North American population consistent with previous analysis reporting a secondary contact between African and North American strains (Duchen *et al.* 2013; Kao *et al.* 2015; Bergland *et al.* 2016) (Figure 3).

**Figure 3.**
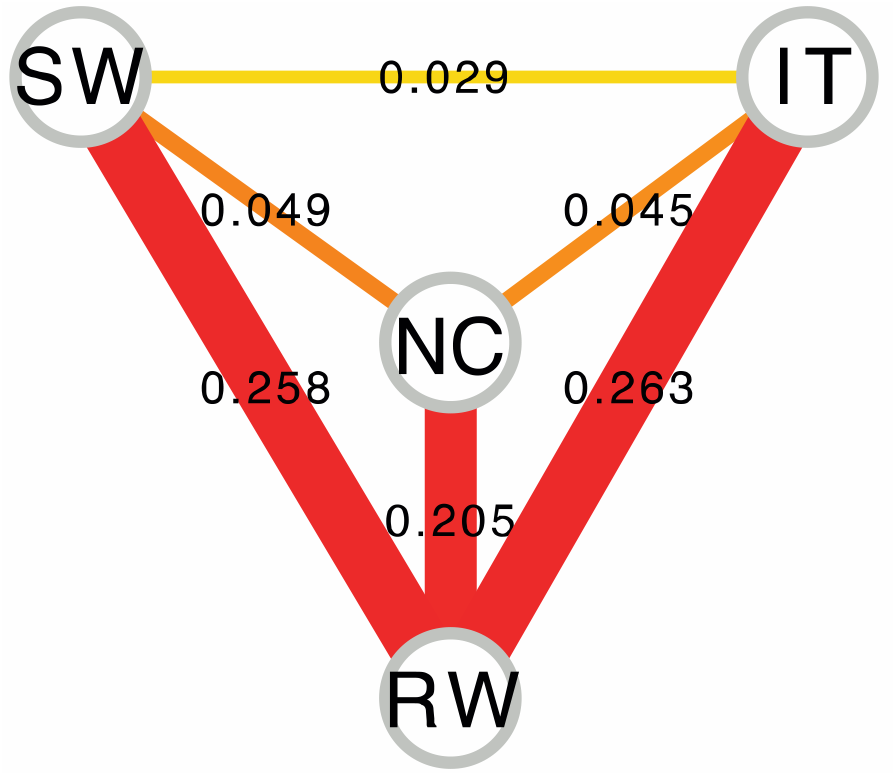
F_ST_ network for the four populations analyzed in this work. Nodes represent the four populations analyzed and edges represent the estimated population distances measured by F_ST_ between all pairs of populations. Increased edge width corresponds with increased differentiation. Spell out acronyms.

Principal component analyses clearly separated strains by continent (Figure 4). The two first principal components (PCs) explained a 39.3% of the observed variance. PC1, which explains 36.3% of the observed variance, separates African strains from non-African strains, and to a lesser extent American and European strains (Figure 4). PC2, which captures 3% of the variance, separates American from European strains. Consistent with F_ST_ estimates, European strains appeared closer to each other than American strains, and African strains (Figure 4). Finally, we also performed model-based global ancestry estimation (Figure 5). The most likely hypothesis according to cross-validation errors, is a model with two or three ancestral populations (Figure S3, Supporting information). The model with two ancestral populations (k = 2) clearly shows a cluster of African ancestry and a cluster of non-African ancestry, with the American population showing an average of 32% of African ancestry. The model with three ancestral populations (k = 3) separates strains by continent (Figure 5). Increasing the number of ancestral populations does not separate the two European populations, suggesting that Swedish and Italian strains belong to a single cluster (Figure 5).

**Figure 4.**
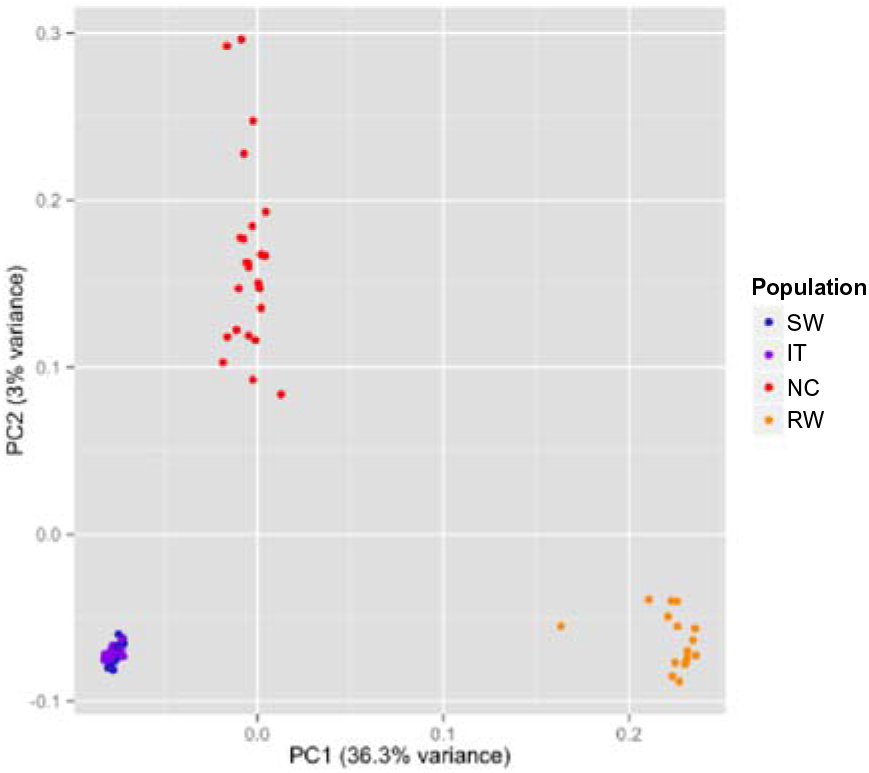
Principal component analysis of the four populations analyzed in this work. Principal component analysis (PCA) was used to visualize the population structure among the 26 Swedish (SW), 16 Italian (IT), 24 North Carolina (NC) and 17 Rwandan (RW) strains. Each point represents one individual strain.

**Figure 5.**
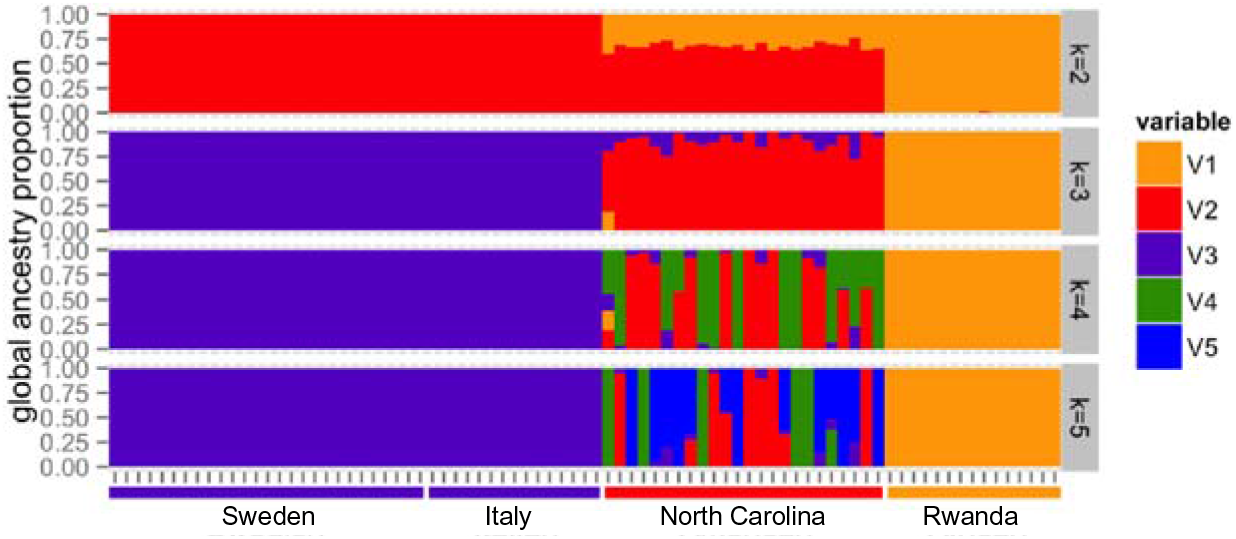
Ancestral population clusters in European, American and African populations. Each bar represents an individual strain and the proportion of the genome that belongs to each ancestral population (k) is filled with a different color.

Overall, as expected, our results are consistent with an out-of-Africa migration, as Rwandan samples are the most genetically diverse (Figure 2). Our results are also consistent with population structure among continents (Figure 3 and 4), and with North American strains showing a considerable proportion of African admixture (Figure 5). Finally, the two European population analyzed showed low levels of population structure (Figure 3 and 4) and no recent African admixture (Figure 5) suggesting that they could be a good dataset to identify loci under spatially varying selection.

### Genomic differentiation between Swedish and Italian populations

To identify signatures of population differentiation between Swedish and Italian strains, we estimated pairwise F_ST_ for all individual SNPs segregating at a minor allele frequency higher than 5% in these two populations (see Material and Methods). The empirical genome-wide distribution of F_ST_, which reflects primarily drift and other non-selective forces, had a maximum around zero and exponentially decays with a long tail of high F_ST_ values (Figure S4A, Supporting information). We found that the levels of differentiation were significantly heterogeneous among chromosomal arms (ANOVA p-value <2e-16; Figure S4B, Supporting information). Therefore, we used the empirical F_ST_ distribution of each chromosomal arm to identify candidate differentiated SNPs in each chromosome. SNPs falling in the top 5%, top 1% or top 0.05% tails of the distribution were considered as candidates (Figure S4C, Supporting information).

To investigate the contribution of inversions to the observed population differentiation patterns, we first estimated the frequency of cosmopolitan inversions in Swedish and Italian populations (Table S4, Supporting information). The only inversion present at a significant frequency was In(2L)t: 18.8% in the Italian strains analysed and 31.5% in the Swedish strains analysed. In(2R)NS was present at low frequencies in both populations, while In(3R)P and In(3L)P were either absent or present at low frequency (Table S4, Supporting information). We thus only considered In(2L)t for the rest of this work.

We then tested whether F_ST_ values inside In(2L)t common cosmopolitan inversions was different from those of the rest of the chromosomal arms. Additionally, we tested whether the proportion of candidate SNPs (top 5%, top 1% and top 0.05%) differed between the inversion and the rest of the chromosomal arm (Figure 6A). The median F_ST_ inside In(2L)t inversion was significantly higher than the median F_ST_ outside this inversion: 0.0080 vs 0.0068 (p-value = 0.0113). However, we did not find a significant accumulation of candidate differentiated loci inside the inversion (Figure 6A).

**Figure 6.**
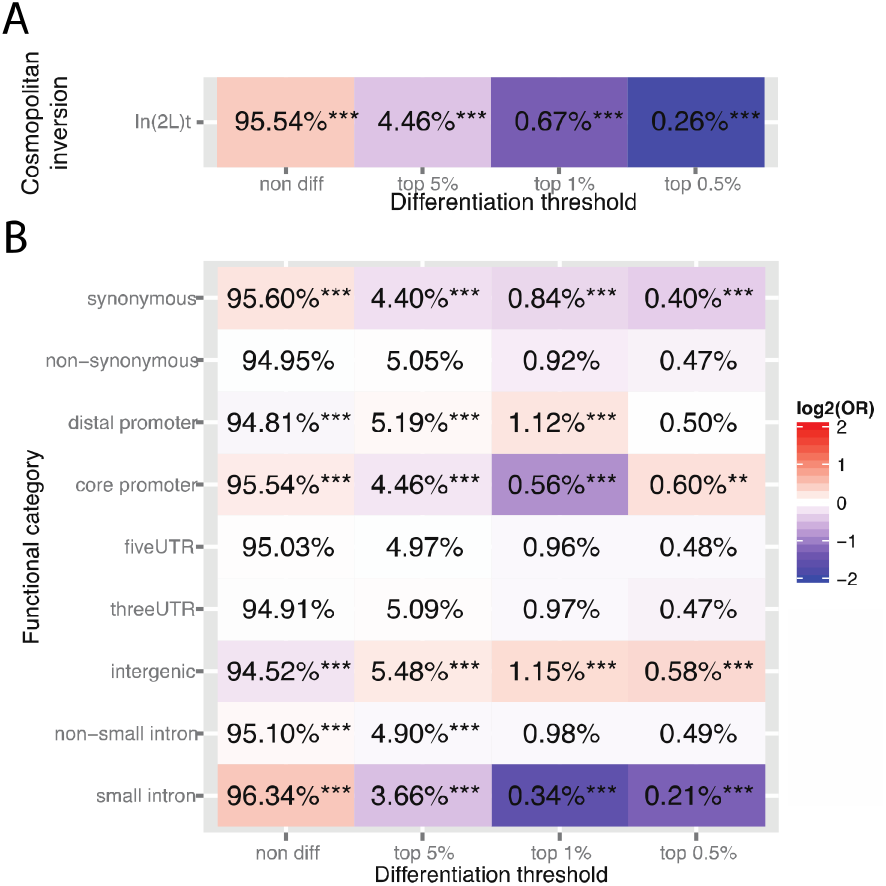
Over- and under-representation of differentiated SNPs in inverted regions and in different functional categories. A) Percentage of non-differentiated and differentiated SNPs (top 5%, top 1% and top 0.5% tails) inside the In(2L)t cosmopolitan inversion. B) Percentage of non-differentiated and differentiated SNPs (top 5%, top 1% and top 0.5% tails) in different functional categories. The odds-ratio is represented as a heat map with red colors indicating over-representation and blue colors under-representation of candidate SNPs. Fisher’s Exact test p-values are indicated as * p < 0.05; ** p < 0.01; and *** p < 0.001.

We then analyzed the functional class of the differentiated SNPs. We first compared the distribution of non-differentiated and differentiated SNPs across the different functional categories and found that as expected, small introns are enriched for non-differentiated SNPs and are depleted of differentiated SNPs (Figure 6B). Our results also showed that core promoter, distal promoter, and intergenic regions are enriched for differentiated SNPs (Figure 6B). Considering small introns as the background and controlling for allele frequency, chromosome, and *In(2L)t* inversion status, we found that although other functional categories are enriched in differentiated SNPs, distal promoter, core promoter and intergenic regions are the three categories that showed a more significant enrichment of candidate SNPs (Table S5, Supporting information).

### Biological processes involved in population differentiation

To identify the biological processes likely to be involved in population differentiation, we first converted the SNP-centric F_ST_ values into gene-centric Z_ST_ scores while controlling for gene length bias (see Material and Methods and Daub *et al.* (2013)). Z_ST_ scores are approximately normally distributed with positive values indicating high levels of population differentiation and negative values indicating low levels of population differentiation (Figure S1, Supporting information). We obtained a Z_ST_ score for 13,140 genes, covering 84% of all annotated *D. melanogaster* genes. We considered as candidate-differentiated genes the 657 genes with a Z_ST_ score in the top 5% of the distribution (Table S6, Supporting information). We then performed GO enrichment analysis using an algorithm based on the classical Fisher’s exact test (*classic*), and three additional more strict algorithms (*elim, weight* and *weight01*) that eliminate local similarities and dependencies between GO terms in order to reduce the false-positive rate (see Material and Methods). The candidate 657 genes were enriched in 43 GO terms according to at least one of the more strict algorithms, which we classified into broader categories to simplify their interpretation (Table S7, Supporting information). The significant biological processes are related to response to xenobiotics, pigmentation, immunity, and developmental processes among others (Table S7, Supporting information). We also checked whether there was overlap between the 43 GO terms identified in this and previous works, and we found that 6 of the GO terms had been previously identified (Table S8, Supporting information). Among the six overlapping GO terms, we found GO:0030707 and GO:0007297, which are related to ovarian follicle cell development and ovarian follicle cell migration, respectively. Interestingly, these processes are associated with diapause in *D. melanogaster* (Saunders *et al.* 1989; Baker & Russell 2009), a complex phenothypic trait involved in the adaptation to temperate regions through overwintering (Schmidt *et al.* 2005; Zonato *et al.* 2017).

### Candidate genes and candidate SNPs involved in population differentiation

To identify the most likely candidate SNPs to play a role in local adaptation, we focused on the 22 genes that belong to the four GO terms that were consistently found to be overrepresented across algorithms: response to insecticide, cuticle pigmentation, neuropeptide signalling pathway, and cell wall macromolecule catabolic process (Table 1). For each gene in these four significant GO terms, we identified the SNP with the highest F_ST_ (Table 1). We checked whether the candidate genes were located inside or nearby the breakpoints of the cosmopolitan inversion In(2L)t. Only one of the 22 candidate genes, *Duox*, was located inside In(2L)t inversion. However, this gene has been previously identified when analysing different natural populations, and also showed expression changes when comparing temperate with tropical populations (Reinhardt *et al.* (2014); Zhao *et al.* (2015); Machado *et al.* (2016)). The strongest F_ST_ signal in the genome corresponds to a coding change in the *Acetylcholinesterase* (*Ace*) gene that is known to confer insecticide resistance and has been previously found to show patterns of population differentiation (Karasov *et al.* 2010; Fabian *et al.* 2012; Reinhardt *et al.* 2014; Machado *et al.* 2016) (Table 1). In this gene, we also identified two additional coding mutations that are known to confer insecticide resistance (3R: 9069408_C/G, Gly303Ala, and 3R: 9063921_C/G, Gly406Ala with F_ST_ values of 0.05 and 0.65, respectively) (Karasov *et al.* 2010). Besides, we also identified two additional coding mutations with unknown functional impact (3R: 9071797_T/C, Thr20Ala; 3R: 9071847_A/G, Ile8Thr with Fst values of 0.3 and 0.38, respectively). A new candidate mutation was also identified for *Cyp6g1* (Table 1). This gene has also been repeatedly reported as a candidate gene in population differentiation studies across continents and it has been experimentally shown to confer resistance to insecticides (Daborn *et al.* 2002; Daborn *et al.* 2007; Battlay *et al.* 2016). Daborn *et al.* (2002) identified a TE insertion upstream of *Cyp6g1* that increased the expression of this gene resulting in resistance to DDT and neonicotinoids (Daborn *et al.* 2002; Daborn *et al.* 2007). More recently, Battlay *et al.* (2016) identified several SNPs in the *Cyp6g1* gene associated with resistance to a different type of insecticide, the organophosphate azinphos-methyl, with the strongest association being from a SNP located in an intron. The mutation identified in this work is a coding mutation, and thus could represent an additional *Cyp6g1* genetic variant involved in insecticide resistance (Table 1). Finally, we found that two other insecticide resistance related genes *Cyp6a8* and *Gr8a* also have coding changes while *Cyp6g2* has a mutation in the core promoter (Table 1).

**Table 1.**
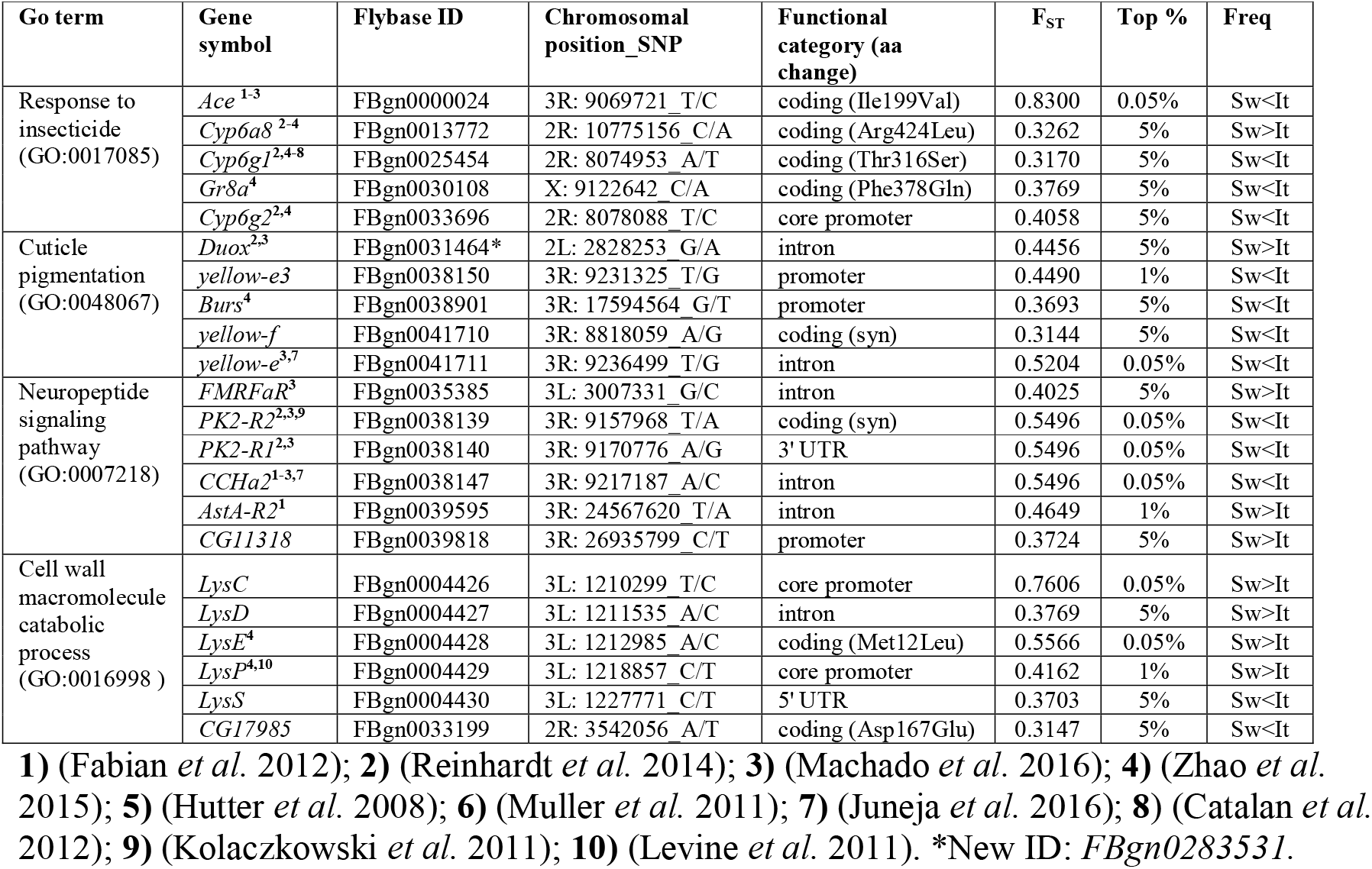
Four GO terms are significantly enriched according to the four algorithms used.

Regarding cuticle pigmentation and neuropeptide signalling, all changes were non-coding changes or synonymous changes, while cell wall macromolecule catabolic process contained non-coding or non-synonymous changes. This later GO term contains the gene with the second highest F_ST_ in the genome: a SNP in the core promoter of *LysC* (Table 1). It is noteworthy that both in cuticle pigmentation (*yellow-e3*, *yellow-e*) and in cell wall macromolecule catabolic process (*LysC*, *LysD*, *LysE*, *LysP*, *LysS*) the differentiated genes are located next to each other, forming two physical clusters of genes that are co-differentiated and functionally related (Table 1). The two clusters are located in regions with high local recombination rates, 2.69 cM/Mbp and 3.15 cM/Mbp respectively, and expand several kilobases, 11.8 kb and 17.8 kb respectively. Thus, although we cannot discard that some of the identified candidate SNPs are linked to the causative adaptive mutation, it could also be that several of them are adaptive (Table 1). Indeed, for *LysP, LysE,* and *yellow-e* there are independent lines of evidence suggesting that they are under selection (Levine *et al.* 2011; Zhao *et al.* 2015; Machado *et al.* 2015, Juneja *et al.* 2016).

We also identified the genes in our dataset that have been reported as candidates in previous studies (Table S1, Supporting information). The number of overlapping genes ranged from zero to 177: while no overlap was found between our candidates and the ones identified by Juneja *et al.* (2016) when comparing two Australian populations, 177 out of 657 genes significantly overlapped between our study and the ones identified by Machado *et al.* (2016) in the North American East coast (Fisher’s exact test, p-value = 0.01904; Table S1, Supporting information). Overall, 363 of the 657 genes (55 %) identified in this study have already been identified in at least one previous study. Interestingly, 35 of these 363 genes, besides showing evidence of population differentiation in other continents, also showed differential gene expression when comparing tropical and temperate populations (Table S9, Supporting information). These 35 genes, which are involved in several ecologically relevant biological processes, are thus good candidates for follow-up functional validation (Table S9, Supporting information).

### Gene set enrichment analysis

Classical GO enrichment analysis is powerful when several genes belonging to the same GO term show strong signals of differentiation that go beyond a threshold (top 5% in this work). However, it fails to detect more subtle trends in which most genes annotated to the same GO term might not belong to the top 5% of the Z_ST_ distribution but as a whole tend to have higher Z_ST_ scores than expected by chance (Subramanian *et al.* 2005). In order to detect those GO terms whose genes tend to occur toward the top of the Z_ST_ distribution, we performed a *Kolmogorov-Smirnov* like test.

A total of 31 GO terms were found to be significantly enriched by the three algorithms and 58 additional GO terms were found to be significantly enriched by at least one of the more strict algorithms. The list of 89 GO terms was classified into broader categories to simplify its interpretation (Table S10, Supporting information). While some broader categories, such as developmental processes and genome organization, overlapped with the GO terms enriched in the top 5% differentiated genes, others such as circadian rhythm were only identified when performing gene set enrichment analysis (Table S10, Supporting information). We found that 13 of the 89 GO terms were previously identified by other works (Table S11, Supporting information).

Overall, the gene set enrichment analyses allowed us to identify additional biological processes potentially involved in differentiation between the two European populations analyzed.

### Transposable elements also contribute to European population differentiation

Besides SNPs, we also investigated whether TEs showed patterns of differentiation between the Swedish and Italian populations. We first genotyped and estimated the frequency of TEs in these two European populations using the *T-lex2* pipeline (Fiston-Lavier *et al.* 2015). We then calculated F_ST_ values for 161 TEs segregating at a minor allele frequency higher than 5% in the two populations (see Material and Methods). The empirical distribution of the F_ST_ estimates for TEs was very similar to the SNPs F_ST_ values distribution (Figure S5, Supporting information). Therefore, we used the SNPs critical F_ST_ values of each chromosomal arm to determine the significantly differentiated TEs. We identified six TEs with significant population differentiation: four of them are present at higher frequencies in the Swedish population while the other two are present at higher frequencies in the Italian population (Table 2).

**Table 2.** Transposable elements showed evidence of population differentiation.

| TE | F<sub>ST</sub> | class | family | Nearby Gene | Name | TE Position | GO (Biological Process) |
| --- | --- | --- | --- | --- | --- | --- | --- |
| FBti0019010 | Sw>It | DNA | pogo | FBgn0033578<sup>1</sup> | BBS4 | Intergenic.<br>1,573bp<br>upstream. | Cilium assembly |
| FBti0019679 | Sw>It | LTR | 1731 | FBgn0031181 | Ir20a | Intergenic.<br>2,487bp<br>downstream. | Detection of chemical<br>stimulus (predicted) |
| FBti0019344 | Sw>It | non-LTR | Rt1a | FBgn0261241<sup>2</sup> | Vps16A | Intergenic.<br>8,616bp<br>downstream. | Syntaxin binding,<br>autophagosome<br>maturation, cellular<br>response to starvation,<br>endosomal transport, eye<br>pigment granule<br>organization |
| FBti0020162 | Sw>It | LTR | Stalker4 | FBgn0062928 | pncr009:3L | Overlapping<br>exon - First<br>non UTR<br>intron 1 | - |
|  |  |  |  | FBgn0052212 | CG32212 | Overlapping<br>5' UTR | - |
|  |  |  |  | FBgn0036874<sup>2</sup> | brv1 | Intergenic.<br>728 bp<br>upstream | Cellular response to cold,<br>detection of temperature<br>stimulus involved in<br>thermoception |
| FBti0019360 | It>Sw | DNA | pogo | FBgn0051358<sup>3</sup> | CG31358 | Intergenic.<br>2,677 bp<br>downstream. | - |
| FBti0020172 | It>Sw | DNA | pogo | FBgn0265959<sup>4</sup> | rdgC | Inside. First<br>non UTR<br>intron 2. | Deactivation of<br>rhodopsin, detection of<br>temperature stimulus<br>involved in sensory<br>perception of pain,<br>phototransduction,<br>protein<br>dephosphorylation,<br>thermotaxis |

Nearby Gene: Genes identified as differentiated between clinal populations: 1) This work; 2)
(Machado *et al.* 2016); 3) (Hutter *et al.* 2008); 4) (Zhao *et al.* 2015).

We identified the genes located nearby these six TEs (< 1kb), and analyzed their associated biological processes GO terms (Table 2). Interestingly, one of the genes nearby insertion *FBti0020162*, which is present at higher frequency in Swedish compared with Italian populations, is involved in cellular response to cold (Table 2). Other ecologically relevant biological processes such as detection of chemical stimulus and response to starvation could be affected by *FBti0019679* and *FBti0019344* insertions respectively (Table 2). Five of the eight genes nearby TEs with significant F_ST_ between Swedish and Italian populations were previously identified in other works (Table 2).

## DISCUSSION

In this work, we tested whether European populations could be a good dataset to disentangle the effects of demography and selection, and thus to identify the genetic basis of adaptation. These two populations were collected in locations with contrasting environments: Stockholm in Sweden, and Castellana Grotte, in Bari, Southern Italy (Figure 1). We first explored the genetic diversity, population structure, and admixture patterns of these two populations in the context of a more global dataset including an African population from the ancestral range of the species, and a North American population (Pool *et al.* 2012; Huang *et al.* 2014). Our results confirmed that there is population structure across continents, and that North American populations show admixture with African populations (Figure 3-5). On the other hand, low levels of population structure and no recent African admixture was detected for the two European populations analyzed (Figure 3-5). Thus, the Italian and Swedish populations analyzed in this work should be a good dataset to identify loci under spatially varying selection.

However, we cannot discard that the European populations analyzed in this work have African admixture from a different source African population. Kao et al (2015) reported some proportion of admixture in strains collected in France when a population from Cameroon was used as a source African population. However, Bergland et al (2015) analyzed admixture in North American and Australian populations using 19 different African populations, including the Gikongoro population used in this study. Admixture was detected independently of the donor population used in the analysis. If there was some degree of admixture that we failed to detect, the patterns of population differentiation identified for some of the European genetic variants could be the result of admixture rather than selection (Duchen *et al.* 2013; Flatt 2015; Kao *et al.* 2015; Bergland *et al.* 2016). To tease apart selective from non-selective signatures of population differentiation, we analyzed our results in the context of all the previous available genome-wide datasets including both genomic and transcriptomic data. Our analyses allowed us to identify new genes likely to be involved in local adaptation, and to provide further evidence for previously identified candidate genes (Table 1 and 2, and Table S6, Supporting information). Thirty-five of the 657 genes identified in this study, besides showing evidence of population differentiation in other continents, also showed evidence of differential gene expression in temperate *vs* tropical populations (Table S10, Supporting information). We argue that these genes, several of which are involved in ecologically relevant processes such as insecticide resistance and pigmentation, are good candidates to underlie local adaptation (Table S10, Supporting information).

For some of the genes already known to play a role in local adaptation, we pinpointed new candidate genetic variants suggesting that different variants might be under selection in different populations. Adding European populations to the already available studies is thus enhancing our understanding of adaptive evolution. This is the case of *Ace* for which, besides providing further evidence for three coding SNPs previously shown to confer resistance to pesticides, we identified two additional coding SNPs that could have also been targets of selection (Karasov *et al.* 2010). Another example is a coding SNP in the *Cyp6g1* gene, which is also involved in insecticide resistance (Daborn *et al.* 2002; Battlay *et al.* 2016). Besides *Ace* and *Cyp6g1*, two other genes functionally annotated as “response to insecticide” were present at higher frequencies in the Italian population. (Table 1). This is consistent with Italian and Swedish populations being subject to different insecticide pressures, which is in agreement with the characteristics of these two populations. While the Italian populations were collected in a vineyard in which pesticides were regularly used, the Swedish populations were collected in urban community gardens and thus are less likely to be subject to strong pesticide use.

Besides response to insecticide, other GO terms such as cuticle pigmentation, defense response, and cell wall macromolecular catabolic process previously related to environmental adaptation were also identified (Table S7, Supporting information). Changes in four of the five genes involved in pigmentation were present at higher frequencies in the Italian population (Table 1). Variation in pigmentation in *D. melanogaster* natural populations has been related to different adaptive phenotypes: adaptation to cold environments, desiccation tolerance, and ultra-violet (UV) resistance (reviewed in True (2003)). The Swedish and Italian populations analyzed in this work differed in temperature, humidity, and UV exposure, and thus it is reasonable to find significant differentiation in SNPs related to pigmentation (Peel *et al.* 2007). Defense response is also considered as an important trait affecting the ability of species to colonize new environments (Early *et al.* 2017). Besides the genes functionally annotated as defense response, three of the six genes functionally annotated as cell wall macromolecule catabolic process have been associated with anti-parasitic defense response (Roxström-Lindquist *et al.* 2004). Finally, the GO analysis also pinpointed six GO terms previously identified in clinal variation studies (Table S8, Supporting information).

Interestingly, two of these terms are associated with ovarian diapause (Saunders *et al.* 1989; Baker & Russell 2009), an ecologically relevant trait putatively involved in the adaptation of *D. melanogaster* to temperate climates (Schmidt *et al.* 2005; Zonato *et al.* 2017).

Besides analyzing individual genes likely to be involved in geographical adaptation, we also performed gene set enrichment analyses. This analysis complements the identification of mutations with strong effects by identifying biological pathways for which several small effect mutations have occurred (Daub *et al.* 2013). We indeed found evidence for polygenic adaptation in several biological processes (Table S10 and Table S11, Supporting information). Some of them, such as circadian rhythm, were not identified with the outlier approach while others, such as detection of chemical stimulus, seem to be affected both by large effect and small effect mutations. These results suggest that in addition to identifying individual genes, gene set enrichment analyses should be performed to get a more complete picture of adaptation.

Finally, while previous studies focused on a single type of genetic variant, besides SNPs we also analyzed TE polymorphisms (Table 2). A total of 4% of the analyzed TEs showed significant patterns of population differentiation. Some of these TEs are located nearby genes involved in ecologically relevant biological processes such as cold adaptation or detection of chemical stimulus (Table 2). Although we analyzed a much bigger dataset compared to previous studies (1,632 TEs vs 902 TEs), the number of TEs for which F_ST_ could be estimated and as such the number of candidate differentiated TEs was relatively small (Gonzalez *et al.* 2008, 2010, 2015). However, our dataset is still incomplete as we focused on the TEs annotated in the reference genome, which is a North American strain (Gonzalez *et al.* 2008). We are thus missing all the TEs present in the two European populations that are not shared with the North American strain sequenced. *De novo* annotation of TEs in the two European populations analyzed in this work would probably result in the identification of more candidate TEs. Thus, further analyses are needed to better assess the role of TEs in local adaptation.

Overall, our results suggest that European populations are a good dataset to advance our knowledge on the genetic basis of environmental adaptation. By analyzing populations with low levels of population structure and no evidence of recent admixture, we were able to confirm and to *de novo* identify genomic targets of spatially variant selection, and to pinpoint genes and biological processes relevant for local adaptation. Understanding adaptation ultimately requires providing functional evidence linking the candidate loci to their relevant phenotypic effects. However, these studies are extremely challenging because they require the identification of the specific phenotype to be examined and the particular experimental conditions in which the phenotype should be tested (Jensen *et al.* 2007). Epistasis and pleiotropy further complicates the establishment of a genotype-phenotype relationship (Lehner 2013; Mackay 2014). Thus, studies as the one reported here that contribute to the identification of a set of candidate genetic variants likely to be under selection are needed in order to pinpoint the most likely candidates for follow-up functional validation. Our study, however, was limited to only two European populations from contrasting environments collected at a single time point. The European continent is very diverse in terms of habitats and climatic areas, and as such, a more complete analysis of different European populations should help identify candidate genes and biological processes under spatially varying selection. Besides spatial variation, analyzing seasonal variation is another venue of research that might be worth exploring in light of recent results obtained in North American populations (Bergland *et al.* 2014, see also Machado *et al.* (2016)).

We conclude that adding European populations to the available genome-wide datasets of *D. melanogaster* populations allowed us to disentangle selective from non-selective signatures of population differentiation in this species. European populations did not show signatures of admixture and showed low levels of population structure and thus are a good dataset to evaluate the contribution of SNPs and TEs to adaptive evolution. The analysis of the genetic variants identified in this work pinpoint genes and biological processes that have been relevant for the adaptation of this species to the out-of-Africa environments. Furthermore, we identified both large-effect and small-effect mutations suggesting that polygenic adaptation has also been relevant in this colonization process.

## Acknowledgements

We thank Anssi Saura and Stefan Escher for helping with the fly collections in Stockholm, (Sweden) and Roberto Torres in Castellana Grotte (Italy). We thank Anna Ullastres and Lain Guio for technical help, and Maite G. Barrón for comments on the manuscript. We also thank Jose Luis Villanueva-Cañas for the estimates of the inversion frequencies. This work was funded by the Spanish Ministry for Economy and Competitivity and FEDER (BFU2011-24397 and BFU2014-57779-P), and by the European Commission (FP7-PEOPLE-2011-CIG-293860 and H2020-ERC-2014-CoG-647900). J.G. was a Ramón y Cajal fellow (RYC-2010-07306).

## Data Accessibility

The datasets generated and analysed during the current study are available in the Sequence Read Archive (SRA) repository, under the BioProject accession: PRJNA390275.

## Author Contributions

JG conceived the study and designed the analysis. LM, GER and JG carried out the analysis, wrote, and revised the manuscript. All authors read and approved the final manuscript.

