## Supplementary figures and images for "Genome-wide patterns of local adaptation in *Drosophila melanogaster*: adding intra European variability to the map"

### Supplementary Materials

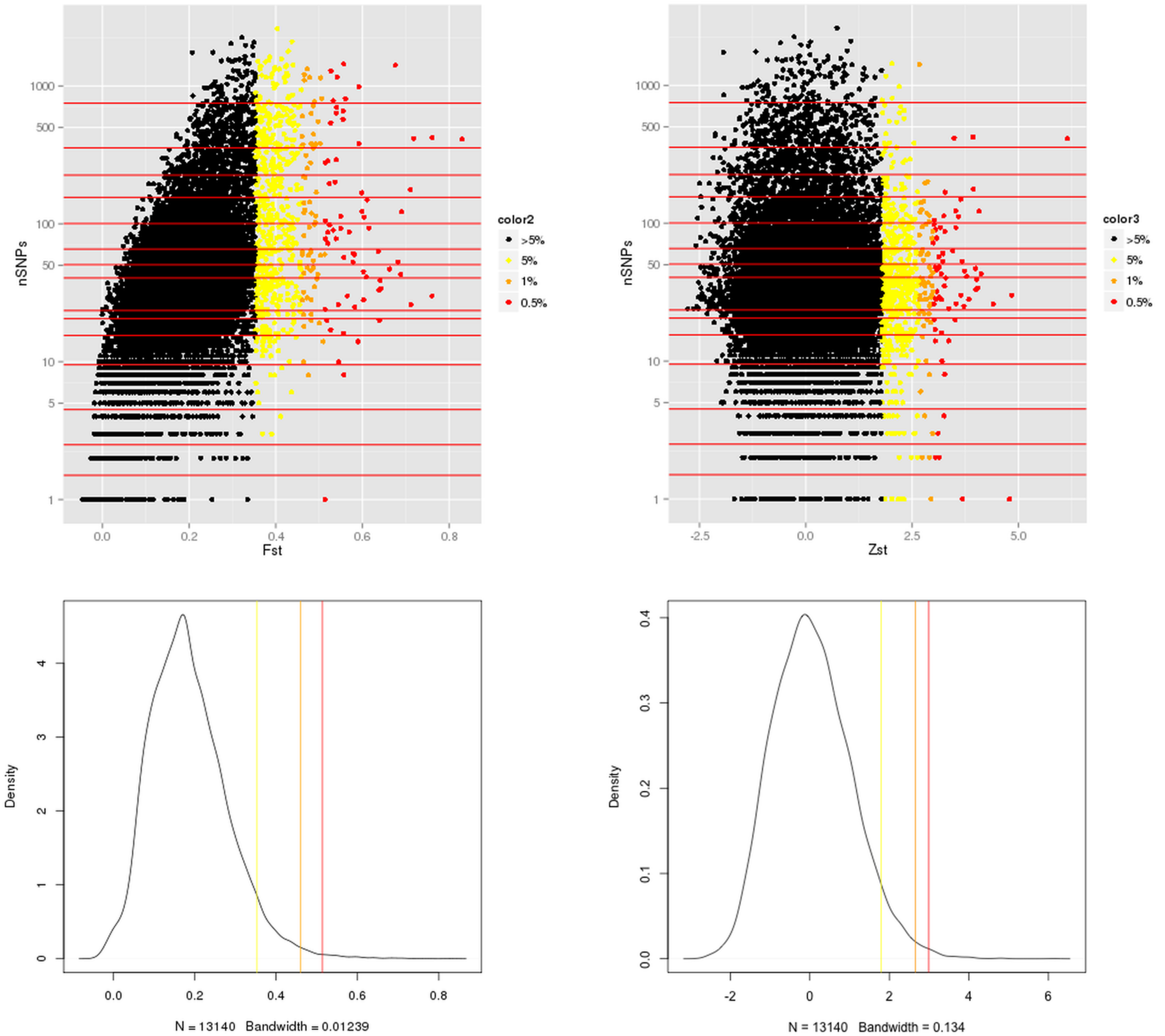

### Supplementary Materials

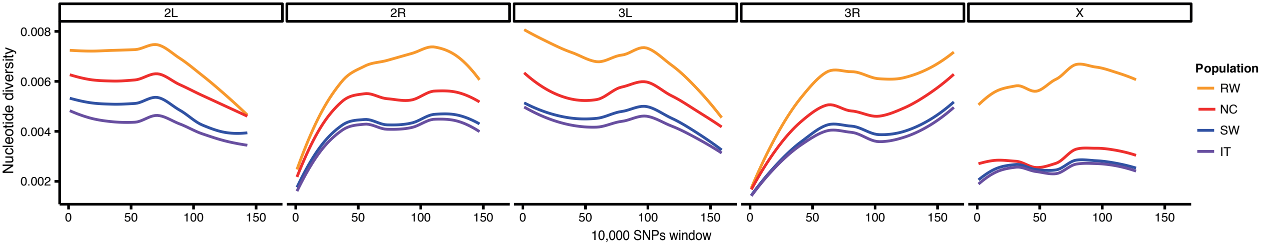

### Supplementary Materials

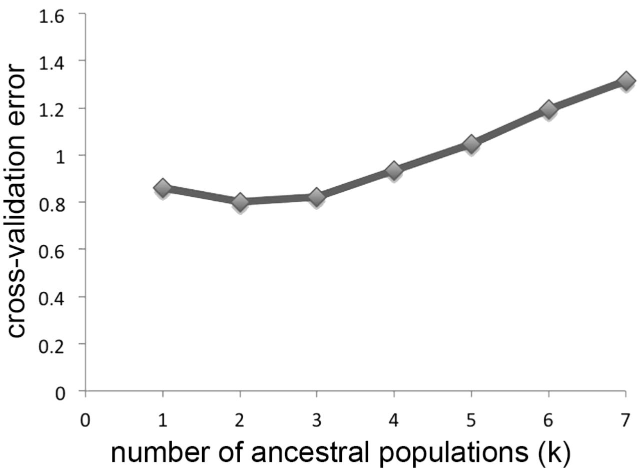

### Supplementary Materials

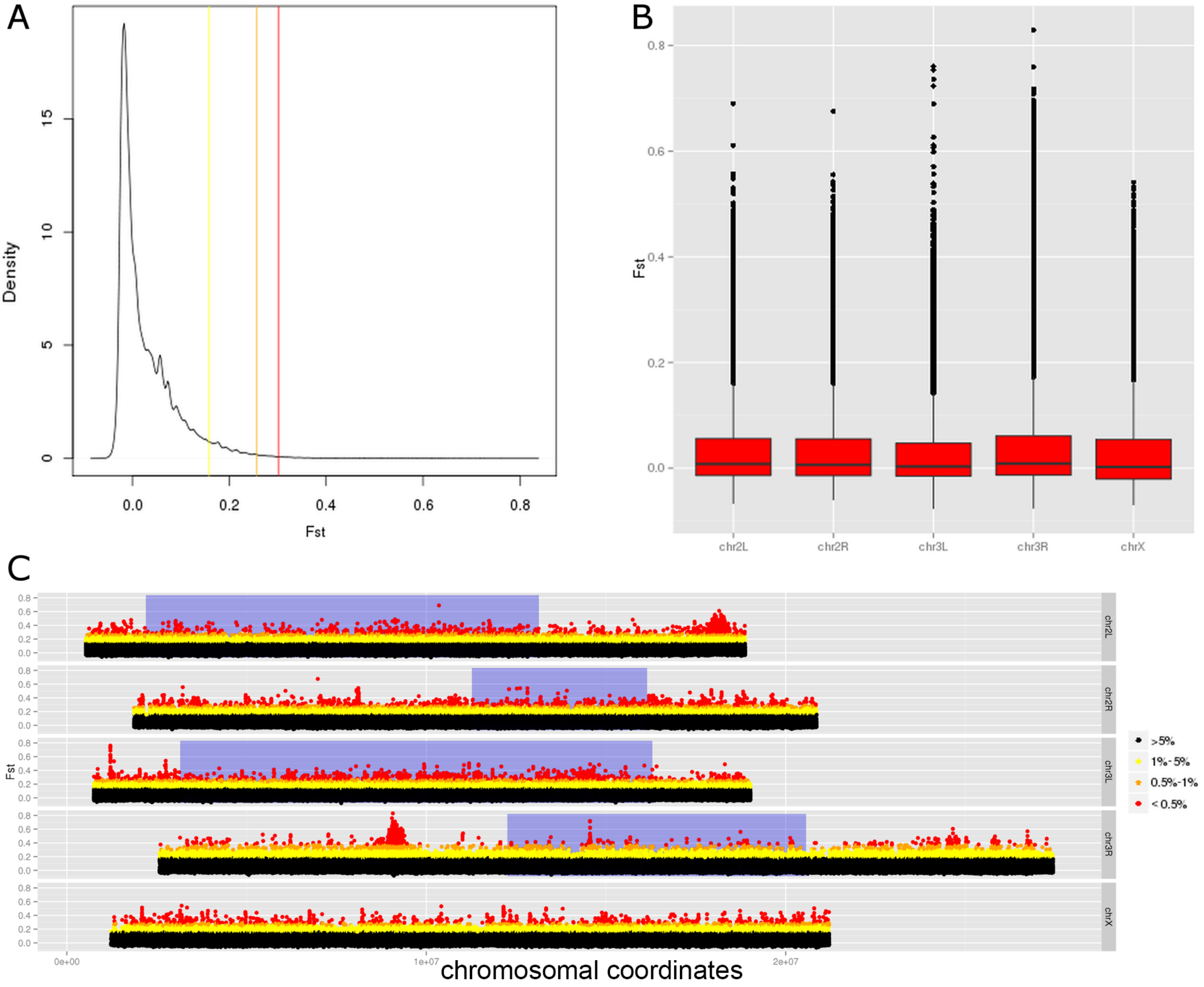

### Supplementary Materials

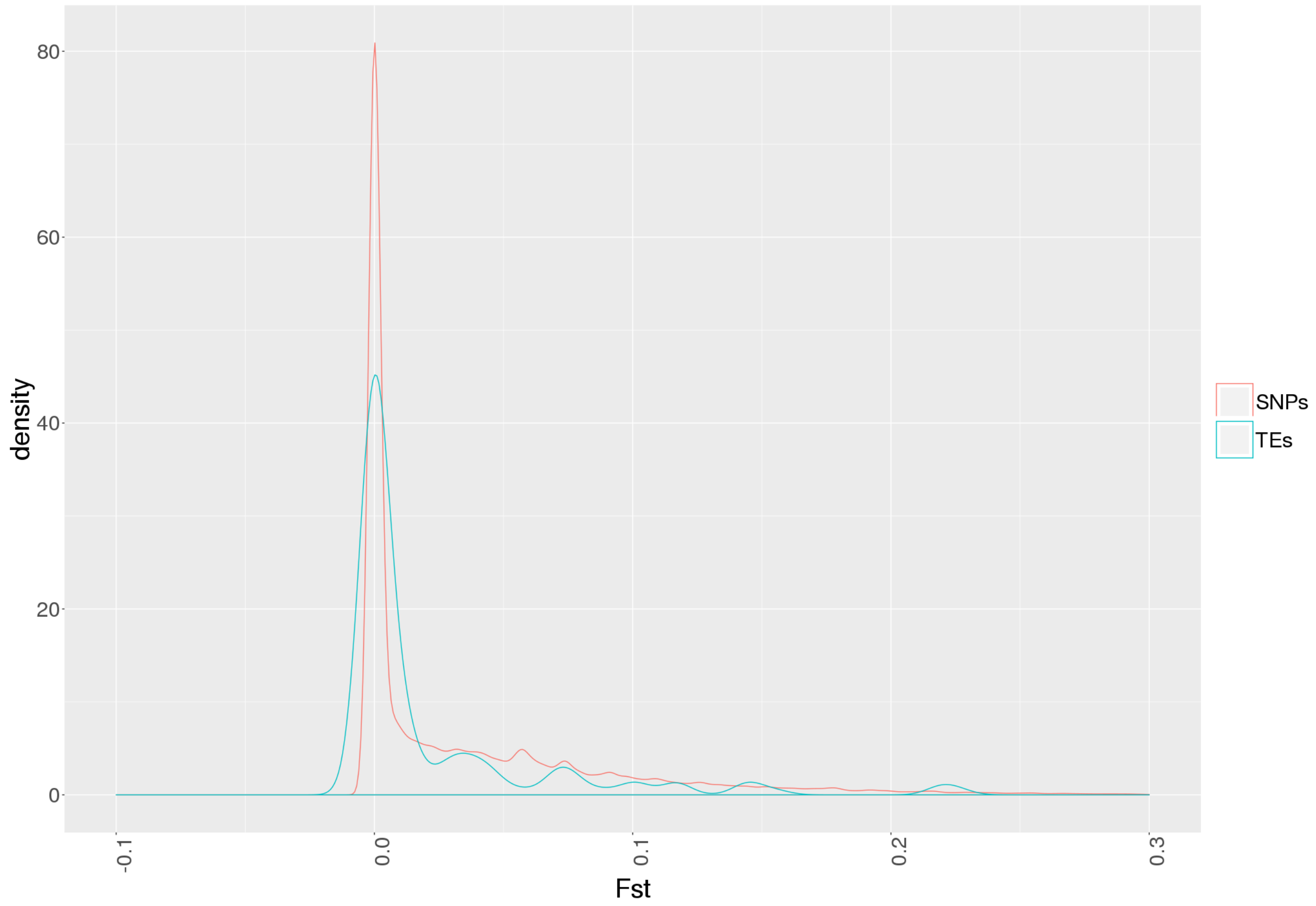
